# Amyloid-beta mediates homeostatic synaptic plasticity

**DOI:** 10.1101/2020.06.22.152066

**Authors:** Christos Galanis, Meike Fellenz, Denise Becker, Charlotte Bold, Stefan F. Lichtenthaler, Ulrike C. Müller, Thomas Deller, Andreas Vlachos

**Author notes:** Department of Neurology, University Hospital of Zurich, Zurich, Switzerland (Denise Becker). Corresponding author, Andreas Vlachos, M.D., Albertstr. 17, 79104 Freiburg, Germany.

## Abstract

The physiological role of the amyloid-precursor protein (APP) is insufficiently understood. Recent work has implicated APP in the regulation of synaptic plasticity. Substantial evidence exists for a role of APP and its secreted ectodomain APPsα in Hebbian plasticity. Here, we addressed the relevance of APP in homeostatic synaptic plasticity using organotypic tissue cultures of *APP^−/−^* mice. In the absence of APP, dentate granule cells failed to strengthen their excitatory synapses homeostatically. Homeostatic plasticity is rescued by amyloid-β (Aβ) and not by APPsα, and it is neither observed in *APP^+/+^* tissue treated with β- or γ-secretase inhibitors nor in synaptopodin-deficient cultures lacking the Ca^2+^-dependent molecular machinery of the spine apparatus. Together, these results suggest a role of APP processing via the amyloidogenic pathway in homeostatic synaptic plasticity, representing a function of relevance for brain physiology as well as for brain states associated with increased Aβ levels.

## INTRODUCTION

In recent years, considerable effort has been directed to better understanding the pathogenic role of APP and its cleavage products in neurodegeneration (Bohm et al., 2015). It has been proposed that the accumulation and deposition of “synaptotoxic” Aβ peptides, which are produced by sequential cleavage of APP by β- and γ-secretase (Lichtenthaler et al., 2011), are responsible for synapse loss, which is regarded as the significant structural hallmark of cognitive decline in Alzheimer’s diseases. In comparison, the physiological functions of cleavage products generated along the “amyloidogenic pathway”, which are also produced in low concentrations in the healthy brain (Cirrito et al., 2003; Seubert et al., 1992; Shoji et al., 1992), are incompletely understood.

Studies employing mouse mutants lacking APP or APP-gene family members (Herms et al., 2004; Magara et al., 1999; Ring et al., 2007; von Koch et al., 1997; Zheng et al., 1995), have shed new light on the role of APP in neuronal migration, synaptogenesis, and synaptic structure and function (Müller et al., 2017). Specifically, alterations in dendritic arborization and spine densities have been reported in pyramidal cells of *APP^−/−^* mice, and these changes are accompanied by defects in the ability of neurons to express long-term potentiation (LTP) of excitatory synaptic strength and alterations in spatial learning [e.g., (Dawson et al., 1999; Li et al., 1996; Muller et al., 1994; Ring et al., 2007; Tyan et al., 2012; Zheng et al., 1995) reviewed in (Ludewig and Korte, 2016; Müller et al., 2017)]. Several of the reported phenotypes of *APP^−/−^* mice are rescued by the APP secreted ectodomain alpha [APPsα, e.g., (Hick et al., 2015; Richter et al., 2018; Tan et al., 2018; Weyer et al., 2014)]. Therefore, it has been proposed that APP could play an important role in neuronal structural and functional plasticity under physiological conditions through the “non-amyloidogenic pathway” which produces APPsα. The physiological function of amyloid-β (Aβ) has, however, remained enigmatic.

In an attempt to learn more about the role of APP and its cleavage products in synaptic plasticity, we here tested for its significance in another major plasticity mechanism (i.e., homeostatic synaptic plasticity). The ability of neurons to adjust their synaptic strength in a compensatory manner is considered to be fundamental for physiological brain function (Styr and Slutsky, 2018; Turrigiano, 1999). It is also a relevant compensatory mechanism in the context of brain diseases and a promising therapeutic target (Andre et al., 2018; Smith-Dijak et al., 2019). In contrast to Hebbian synaptic plasticity, e.g., LTP, homeostatic plasticity is based on negative feedback mechanisms (Lisman, 2017; Turrigiano, 2008). Meanwhile, several molecular players have been identified that control homeostatic synaptic plasticity (Cingolani et al., 2008; Goddard et al., 2007; Stellwagen and Malenka, 2006; Sun and Turrigiano, 2011; Walters and Josselyn, 2019). Recently, a potential role of APP in homeostatic synaptic plasticity has been discussed (Andre et al., 2018; Hoe et al., 2012; Jang and Chung, 2016; Styr and Slutsky, 2018). However, the mechanistic link between APP and homeostatic plasticity remains unclear because homeostatic plasticity could be induced by APP and its cleavage products or indirectly by synapse loss and cell death [i.e., denervation-induced homeostatic adaptation; e.g., (Deller and Frotscher, 1997; Steward, 1994; Vlachos et al., 2012; Vlachos et al., 2013)]. Here, we studied the role of APP in non-diseased brain tissue and report an essential role of Aβ in homeostatic plasticity of excitatory neurotransmission, suggesting that this could be one of the major physiological functions of Aβ in the normal brain.

## RESULTS

### Homeostatic synaptic plasticity is not observed in dentate granule cells of APP-deficient entorhinal-hippocampal tissue cultures

Considering the role of the hippocampal formation and specifically the dentate gyrus in memory formation (Aimone et al., 2011; Friedman and Goldman-Rakic, 1988), 3-week-old (?18 days in vitro; div) organotypic tissue cultures containing the entorhinal cortex and the hippocampus were prepared from *APP^+/+^* and *APP^−/−^* mice–including age- and time-matched *APP^+/+^* littermates obtained from *APP*^+/-^ intercrossing (Figure 1*A, B*). Tissue cultures were treated with tetrodotoxin (TTX; 2 μM; 2 days) to induce homeostatic synaptic plasticity, and α-amino-3-hydroxy-5-methyl-4-isoxazolepropionic acid (AMPA) receptor-mediated miniature excitatory postsynaptic currents (mEPSCs) were recorded from individual dentate granule cells (Figure 1*C*) to assess compensatory (i.e., homeostatic) synaptic changes.

**Figure 1:**
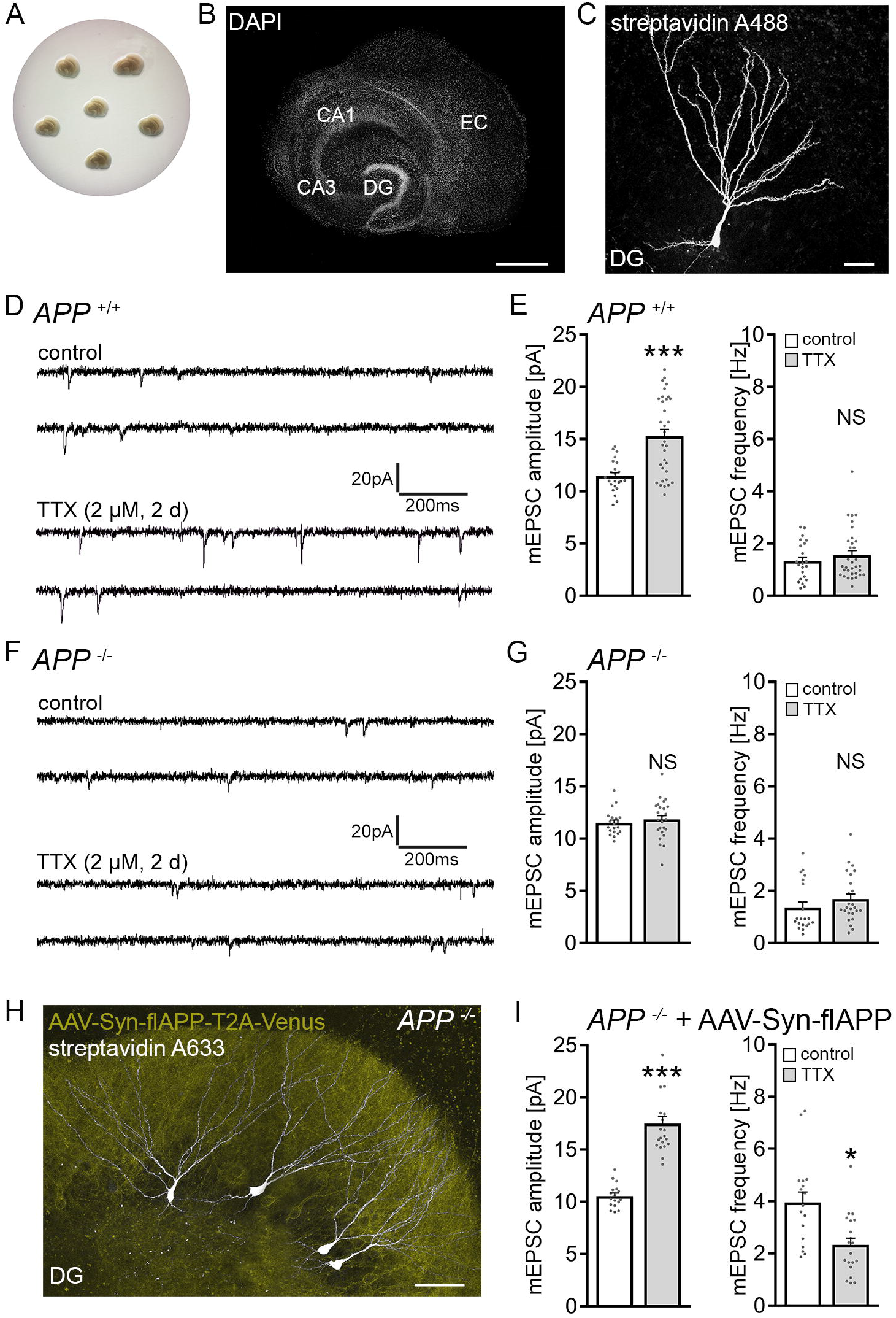
Dentate granule cells of APP-deficient entorhinal-hippocampal tissue cultures do not express homeostatic synaptic plasticity. **(A, B)** Example of three-week old entorhinal-hippocampal tissue cultures on a membrane insert and overview at higher magnification of a representative culture. DAPI nuclear staining used for visualization of cytoarchitecture (DG, dentate gyrus; EC, entorhinal cortex; CA1 and CA3, Cornu Ammonis areas 1 and 3). Scale bar 500 μm. **(C)** Patched dentate granule cell filled with biocytin and identified post-hoc with streptavidin A488. Scale bar 25 μm. **(D, E)** Sample traces and group data of AMPA receptor mediated miniature excitatory postsynaptic currents (mEPSCs) recorded from granule cells in vehicle-treated (control) and tetrodotoxin (TTX)-treated *APP^−/−^* cultures (control, n = 23 cells from 6 cultures; TTX, n = 33 cells from 8 cultures; Mann-Whitney test). **(F, G)** Sample traces and group data of APP-deficient (*APP^−/−^*) dentate granule cells (control, n = 20 cells from 6 cultures; TTX, n = 25 cells from 7 cultures; Mann-Whitney test). **(H)** Example of post-hoc identified recorded dentate granule cells (streptavidin 633, white) in an *APP* tissue culture transduced with adeno-associated viral vectors expressing full length *APP^−/−^* (AAV-flAPP Venus; yellow). Scale bar 50 μm. **(I)** Expression of the flAPP rescues the ability of dentate granule cells in *APP^−/−^* tissue cultures to express TTX-induced homeostatic synaptic plasticity (control, n = 17 cells from 6 cultures; TTX, n = 20 cells from 7 cultures; Mann-Whitney test). Individual data points are indicated in this and the following figures by grey dots. Values represent mean ± s.e.m. (* p < 0.05, *** p < 0.001; NS, no significant difference).

In line with previous work [e.g., (Echegoyen et al., 2007; Kim and Tsien, 2008; Strehl et al., 2018; Turrigiano et al., 1998; Vlachos et al., 2013)], a homeostatic increase in excitatory synaptic strength (i.e., a robust increase in mEPSC amplitudes) was observed in the wild-type tissue cultures (Figure 1*D, E*). In *APP^−/−^* preparations, no significant changes in mEPSC properties were observed in dentate granule cells (Figure 1*F, G*). Specifically, mean mEPSC amplitude was 11.5 ± 0.3 pA in vehicle-only-treated and 11.8 ± 0.4 pA TTX-treated *APP^−/−^* dentate granule cells (p = 0.4; Mann-Whitney-test).

In an attempt to rescue the ability of granule cells to express homeostatic synaptic plasticity, *APP^−/−^* tissue cultures were transfected with a bicistronic adeno-associated viral vector expressing full-length murine APP (flAPP) and membrane-anchored Venus linked by a T2A site under the control of the neuronal synapsin promoter (AAV-Syn-flAPP-T2A-Venus; Figure 1*H*). Cultures were transduced at 4-5 div and allowed to mature for at least 18 div before experimental assessment. A significant compensatory increase in mEPSC amplitudes from 10.5 ± 0.3 pA to 17.5 ± 0.7 pA (p < 0.001; Mann-Whitney-test; Figure 1*I*) was observed in the TTX-group, while increased mEPSC frequencies were observed in the untreated cultures reduced after TTX treatment (Figure 1*I*). We conclude from these results that postnatal expression of APP is required for TTX-induced homeostatic scaling of excitatory synapses to occur in cultured dentate granule cells.

Based on our recent work (Lenz et al., 2019), we also tested for changes in inhibitory synaptic strength, and we did not detect TTX-induced changes in miniature inhibitory postsynaptic currents (mIPSCs) of dentate granule cells, either in *APP^+/+^* or *APP^−/−^* tissue cultures (Figure S1). Hence, we focused on the role of APP in excitatory synaptic scaling.

### No significant alterations in basic functional and structural properties of APP-deficient dentate granule cells

To test whether alterations in baseline synaptic activity explain the inability of *APP^−/−^* granule cells to express homeostatic excitatory synaptic plasticity, spontaneous excitatory and inhibitory postsynaptic currents were recorded in a different set of 3-week-old tissue cultures (Figure 2*A–D*). No significant differences between the two genotypes were observed in these experiments. Similarly, the input-output properties of dentate granule cells (Figure 2*E, F*), as well as basic properties of action potentials (Figure 2*G*), were not significantly different between the two groups.

**Figure 2:**
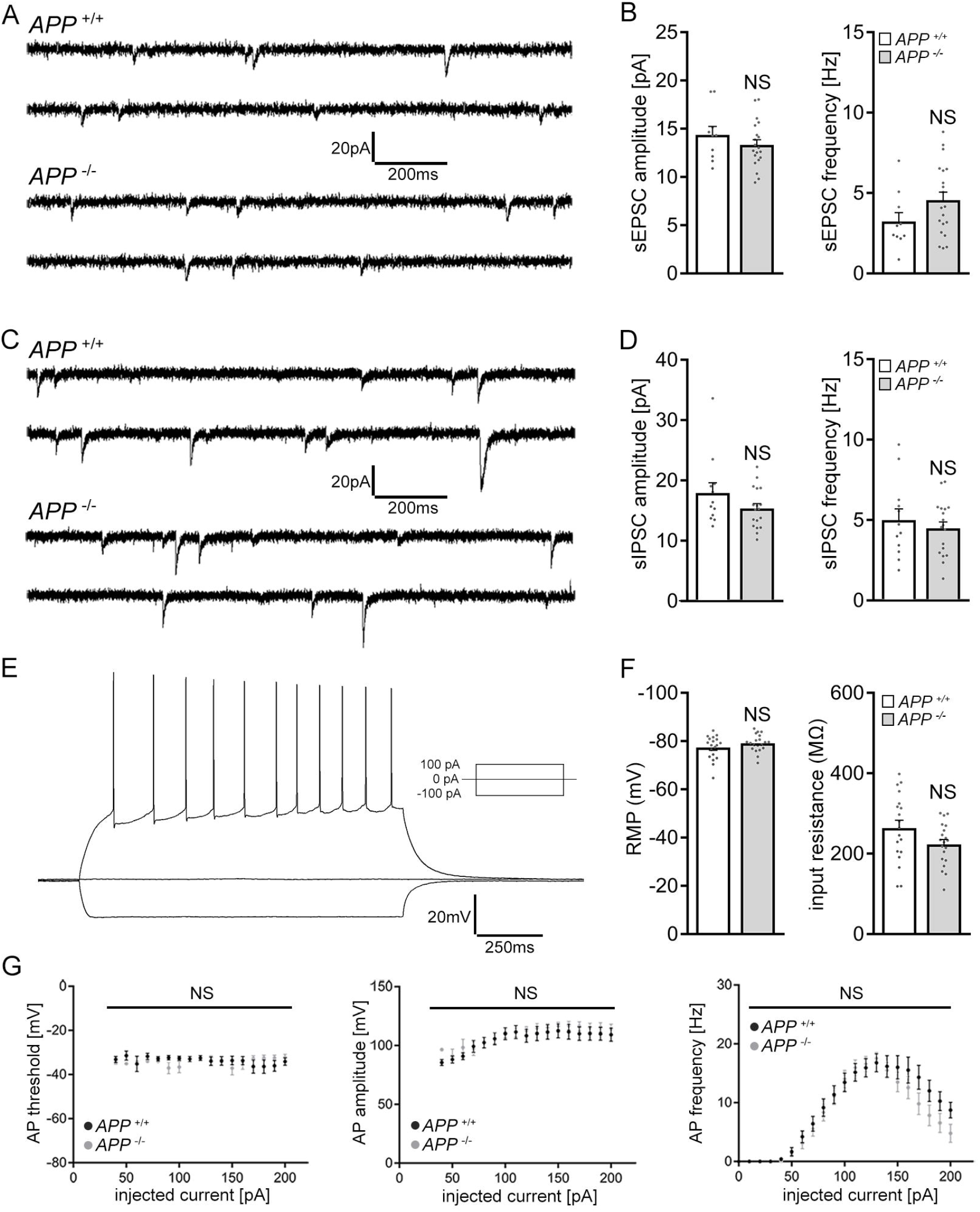
APP deficiency does not affect basic functional properties of dentate granule cells. **(A, B)** Sample traces and group data of spontaneous excitatory postsynaptic currents (sEPSC) recorded from dentate granule cells of *APP^+/+^* and *APP^−/−^* tissue cultures (*APP^+/+^*, n = 10 cells from 4 cultures, *APP^−/−^*, n = 20 cells from 6 cultures; Mann-Whitney). **(C, D)** Sample traces and group data of spontaneous inhibitory postsynaptic currents (sIPSC) from granule cells of *APP^+/+^* and *APP^−/−^* tissue cultures (*APP^+/+^*, n = 12 cells from 4 cultures, *APP^−/−^*, n = 18 cells from 6 cultures; Mann-Whitney test). **(E-G)** Sample traces and group data for input-output properties of dentate granule cells of *APP^+/+^* and *APP^−/−^* tissue cultures. (RMP, resting membrane potential, AP, action potential; *APP^+/+^*, n = 19 cells from 5 cultures, *APP^−/−^*, n = 20 cells from 5 cultures; NS; Mann-Whitney test and 2way ANOVA). Values represent mean ± s.e.m. (NS, no significant difference).

We also tested for variations in basic structural properties of granule cells between *APP^+/+^* and *APP^−/−^* preparations. As shown in Figure S2, an assessment of total dendritic branch length and Sholl analysis revealed no significant differences between the genotypes (Figure 3*A–C*). We also did not find any significant differences in spine densities (Figure 3*D–F*), and synapses were regularly observed in electron microscopy cross-sections of recorded and *post-hoc* stained *APP^−/−^* dentate granule cells (Figure 3*F*). Hence, basic functional and structural alterations do not readily explain the inability of *APP^−/−^* granule cells to scale their excitatory synapses.

**Figure 3:**
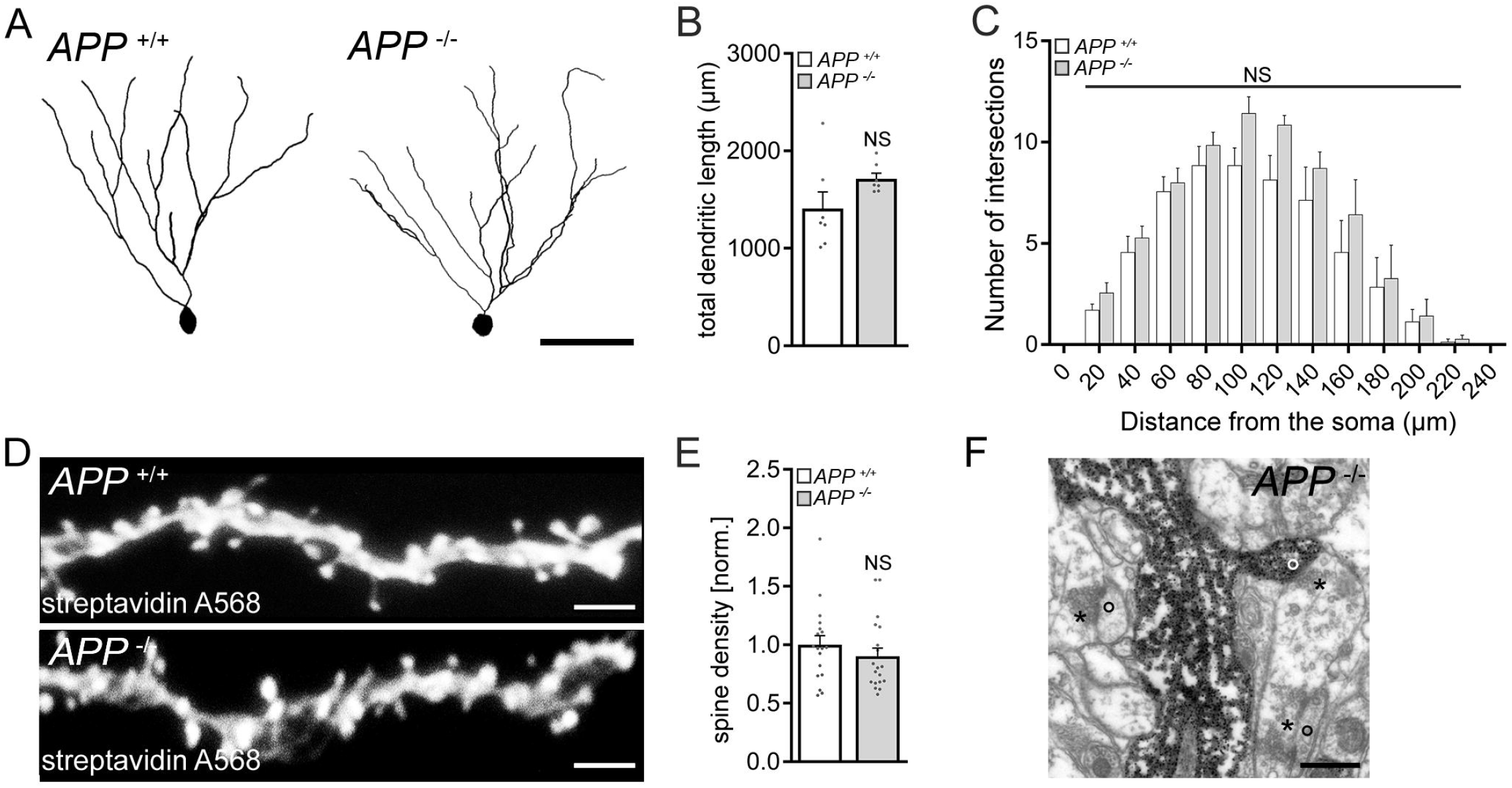
No significant structural differences are observed in granule cells of APP-deficient preparations. **(A – C)** Examples and group data of total dendritic length and Sholl sphere analysis of dentate granule cells in *APP^+/+^* and *APP^−/−^* tissue cultures (n = 7 cells from 7 cultures per group; NS; Mann-Whitney test and 2way ANOVA), scale bar 50 μm **(D, E)** Spine density analysis of distal dendritic segments of dentate granule cells in the outer molecular layer (OML; n = 18 – 19 segments from 6 tissue cultures per group; Mann-Whitney test). Scale bar 5 μm. **(F)** Electron micrograph of synaptic contacts in the OML of *APP^−/−^* tissue cultures. Examples of presynaptic compartments indicated by asterisks, postsynapses by circles. Scale bar 0.5 μm. Values represent mean ± s.e.m. (NS, no significant difference).

### A secreted factor rescues the ability of APP-deficient dentate granule cells to express homeostatic synaptic plasticity

In our viral transduction experiments, in which we used flAPP to rescue homeostatic synaptic plasticity of *APP^−/−^* granule cells (c.f., Figure 1*H, I*), we noticed that granule cells not expressing flAPP also showed increased mEPSC amplitudes after TTX treatment. Thus, we hypothesized that a secreted factor–and not flAPP expression *per se*–mediates the effects of APP on TTX-induced homeostatic synaptic plasticity.

To test this hypothesis, *APP^−/−^* tissue was cultured with 5 *APP^+/+^* cultures on the same membrane insert as shown in Figure 4*A*. In these experiments, a significant increase in excitatory synaptic strength was detected in dentate granule cells of TTX-treated *APP^−/−^* cultures (Figure 4*B, C*). We conclude that a secreted molecular signal originating from *APP^+/+^* tissue is sufficient to rescue the ability of granule cells in the *APP^−/−^* cultures to express TTX-induced homeostatic synaptic plasticity. The results of these experiments also indicate that the presence of APP in the target region is not required (i.e., APP *per se* does not act as a receptor for signaling pathways relevant for TTX-induced excitatory synaptic scaling).

**Figure 4:**
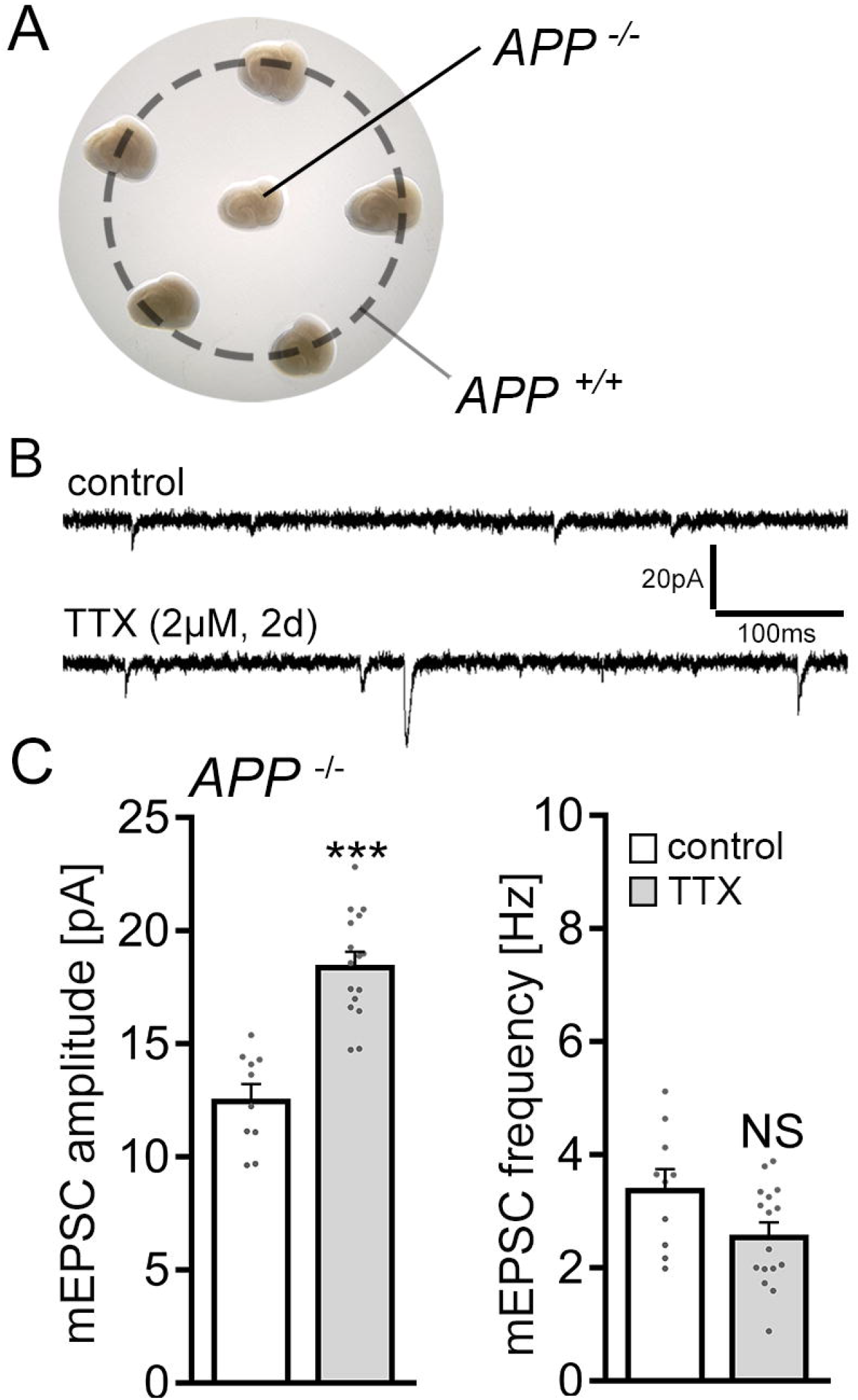
A secreted factor mediates the ability of APP−/− dentate gyrus granule cells to express homeostatic synaptic plasticity. **(A)** Example of co-cultured *APP^+/+^* and *APP^−/−^* tissue preparations on the same membrane insert. **(B, C)** Sample traces and group data of AMPA receptor mediated miniature excitatory postsynaptic currents (mEPSCs) recorded from dentate gyrus granule cells in vehicle-treated (control) and tetrodotoxin (TTX)-treated (2 μM, 2 d) *APP^−/−^* tissue cultures (control, n = 10 cells from 4 cultures; TTX, n = 16 cells from 5 cultures; Mann-Whitney test). Values represent mean ± s.e.m. (*** p < 0.001; NS, no significant difference).

### APPsα does not rescue homeostatic synaptic plasticity in APP-deficient preparations

Previous work revealed that APPsα rescues several phenotypes observed in *APP^−/−^* mice, including alterations in Hebbian synaptic plasticity [i.e., LTP (e.g., (Fol et al., 2016; Hick et al., 2015; Richter et al., 2018; Ring et al., 2007)]. We, therefore, tested whether APPsα could be the secreted factor that rescues the ability of *APP^−/−^* granule cells to express homeostatic synaptic plasticity in our experiments (c.f., Figure 4).

First, *APP^−/−^* cultures were treated with APPsα at a concentration that rescues LTP in acute hippocampal slices [10 nM; (Hick et al., 2015)], and TTX-induced synaptic scaling was probed. In these experiments no homeostatic synaptic adjustment was observed (Figure 5*A, B*), thus confirming once more our major finding (i.e., alterations in homeostatic synaptic plasticity of *APP^−/−^* preparations).

**Figure 5:**
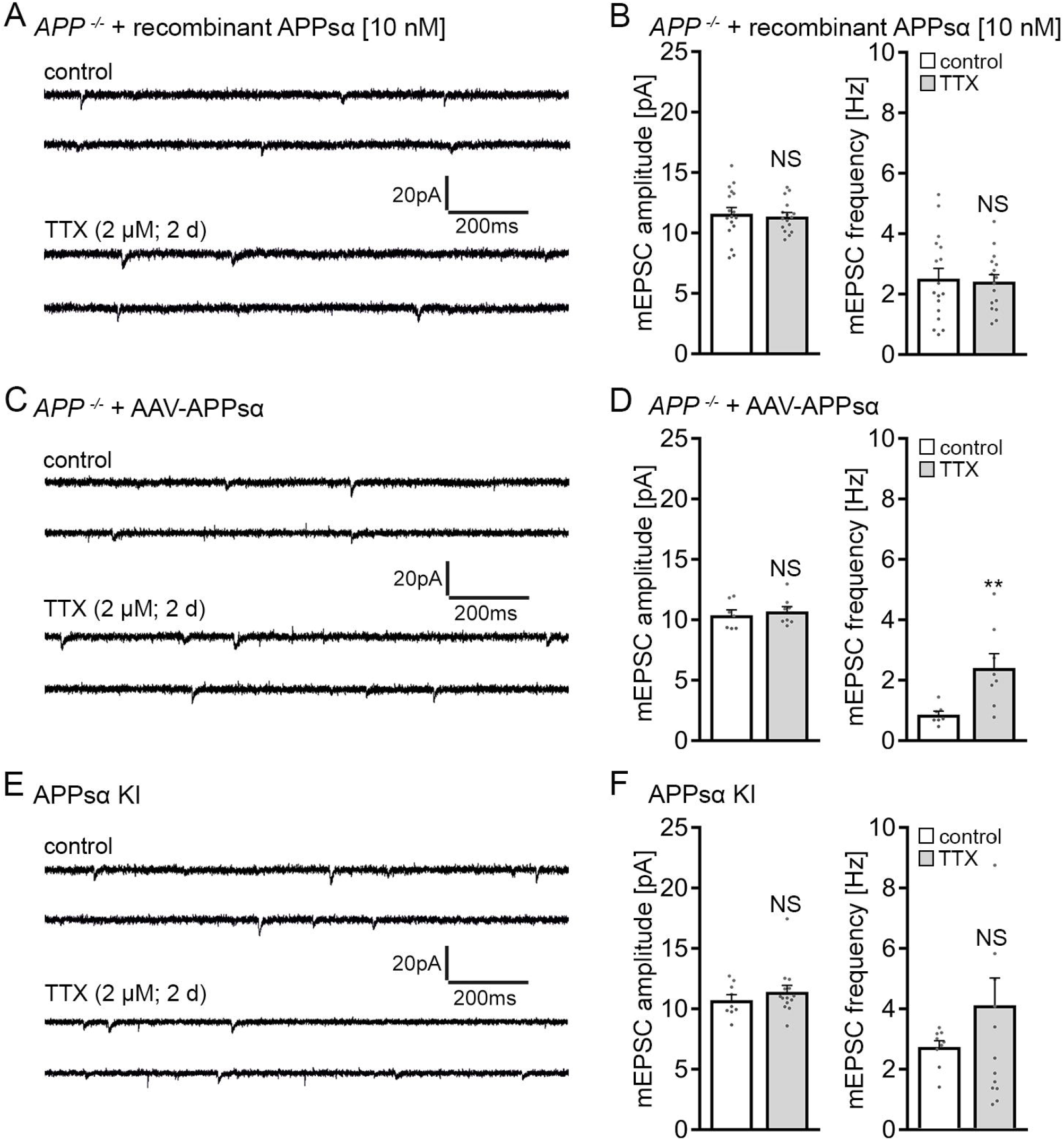
APPsα does not rescue the ability of APP-deficient dentate granule cells to express homeostatic synaptic plasticity. **(A, B)** Sample traces and group data of AMPA receptor mediated miniature excitatory postsynaptic currents (mEPSCs) recorded from granule cells in vehicle-treated (control) and tetrodotoxin (TTX)-treated (2 μM, 2 d) APP^−/−^ tissue cultures in the presence of recombinant APPsα (10 nM; control, n = 17 cells from 5 cultures; TTX, n =16 cells from 5 cultures; Mann-Whitney test). **(C, D)** Sample traces and group data of AMPA receptor mediated mEPSCs from granule cells in APP−/− tissue cultures transduced with adeno-associated viral vectors expressing APPsα (control, n = 7 cells; TTX, n = 8 cells; from 2 cultures per group; Mann-Whitney test). **(E, F)** Sample traces and group data of AMPA receptor mediated mEPSCs from granule cells in tissue cultures prepared from APPsα knock-in (KI) mice (control, n = 9 cells from 3 cultures; TTX, n = 14 cells from 4 cultures; Mann-Whitney test; for mEPSC frequency one data point is outside the axis limits in the TTX group). Values represent mean ± s.e.m. (** p < 0.01; NS, no significant difference).

We next resorted to adeno-associated viral transduction of secreted APPsα (AAV-Syn-T2A-Venus-APPsα) using the same protocol as described for flAPP (c.f., Figure 1*H, I*). The expression of this viral construct was previously employed to successfully rescue LTP-defects in *APP^−/−^* mice (Fol et al., 2016). Again, in our experimental setting no compensatory increase in mean mEPSC amplitude was observed after TTX treatment as compared to age- and time-matched vehicle-only-treated APPsα transfected *APP^−/−^* tissue cultures (Figure 5*C, D*). Interestingly, an increase in mEPSC frequencies back to baseline was detected in the TTX-group in these experiments (Figure 5*D*). Notably, these experiments also indicated that viral transduction *per se* does not rescue the ability of *APP^−/−^* granule cells to express homeostatic synaptic plasticity (c.f., Figure 1*H, I*).

Finally, tissue cultures from *APPsα-KI* mice were prepared (Ring et al., 2007), which express APPsα constitutively while lacking transmembrane APP and Aβ; this represents another approach for rescuing LTP (Ring et al., 2007). Because in these experiments we also did not observe homeostatic plasticity (Figure 5*E, F*), we conclude that APPsα does not rescue TTX-induced homeostatic synaptic plasticity of dentate granule cells in APP^−/−^ tissue cultures, at least not by using the above-described experimental approaches that all rescue LTP.

### Scavenging endogenous APPsα with a specific antibody does not block homeostatic synaptic plasticity

We next tested for the effects of endogenous APPsα by treating wild-type tissue cultures with TTX (2 μM; 2 days) in the presence of a specific antibody that binds APPsα [i.e., the N-terminal APP-E1 domain; JRD32; 1.3 μg/ml; (Hick et al., 2015)]. As shown in Figure 6*A*, a compensatory increase in mEPSC amplitudes was observed in these experiments, thus providing additional evidence that APPsα is not involved in mediating homeostatic synaptic plasticity.

**Figure 6:**
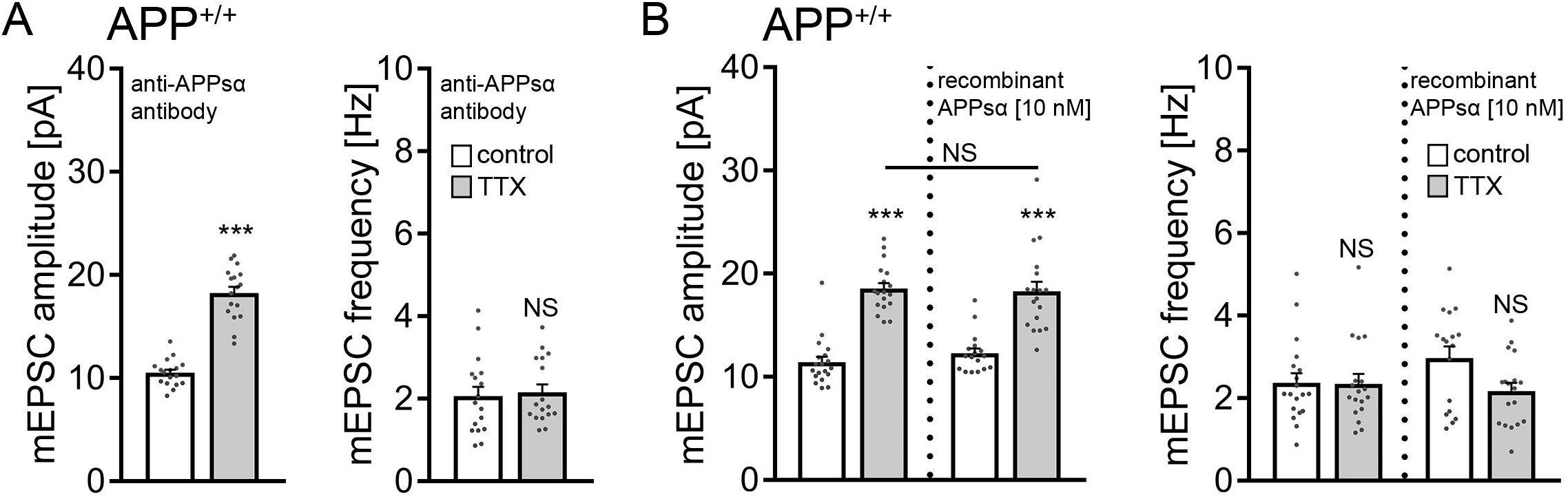
APPsα has no apparent effect in TTX-induced homeostatic plasticity. **(A)** Group data of AMPA receptor mediated miniature excitatory postsynaptic currents (mEPSC) recorded from granule cells in vehicle-treated (control) and tetrodotoxin (TTX)-treated (2 μM, 2 d) *APP^+/+^* tissue cultures in the presence of a specific anti-APPsα antibody (JRD32; 1.3 μg/ml; n = 17 cells from 6 cultures per group; Mann-Whitney test). **(B)** Recombinant APPsα (10 nM) does not affect the TTX-induced increase in mEPSC amplitudes of *APP^+/+^* dentate granule cells (control, n = 19 cells from 7 cultures; TTX, n = 18 cells from 6 cultures; APPsα, n = 17 cells from 6 cultures; TTX + APPsα, n = 18 cells from 6 cultures; Kruskal-Wallis-test followed by Dunn’s post-hoc test). Values represent mean ± s.e.m. (*** p < 0.001; NS, no significant difference).

Because treatment with recombinant APPsα (10 nM) also did not cause aberrant “over”-scaling in *APP^+/+^* dentate granule cells (Figure 6*B*), together with the experiments carried out in *APP^−/−^* cultures (c.f., Figure 5), we are confident to conclude that APPsα is not a major regulator of homeostatic synaptic plasticity (this study) whereas it does promote LTP [e.g., (Fol et al., 2016; Hick et al., 2015; Richter et al., 2018; Ring et al., 2007)].

### Aβ rescues homeostatic synaptic plasticity in APP-deficient preparations

We then considered the amyloidogenic processing pathway and the potential for Aβ involvement in mediating homeostatic synaptic plasticity. Another set of *APP^−/−^* tissue cultures was treated for 2 days with TTX and with a synthetic Aβ protein fragment 1 – 42 [1.5 μM; (Novotny et al., 2016)]. Notably, a full-sized homeostatic synaptic scaling response was observed in these experiments, whereas 2 days of Aβ_1-42_ treatment had no effect on baseline mEPSC amplitudes and frequencies (Figure 7*A*): in the presence of Aβ_1-42_, mEPSC amplitudes increased from 10.9 ± 0.5 pA to 16.8 ± 0.9 pA after 2 d TTX treatment (p < 0.001; Kruskal-Wallis-test followed by Dunn’s post-hoc test). Because similar experiments with the synthetic Aβ protein fragment 42 – 1 (1.5 μM) did not show any significant changes in mEPSC properties (Figure 7*B*), we conclude that the application of exogenous Aβ_1-42_ rescues TTX-induced homeostatic synaptic plasticity in *APP^−/−^* preparations.

**Figure 7:**
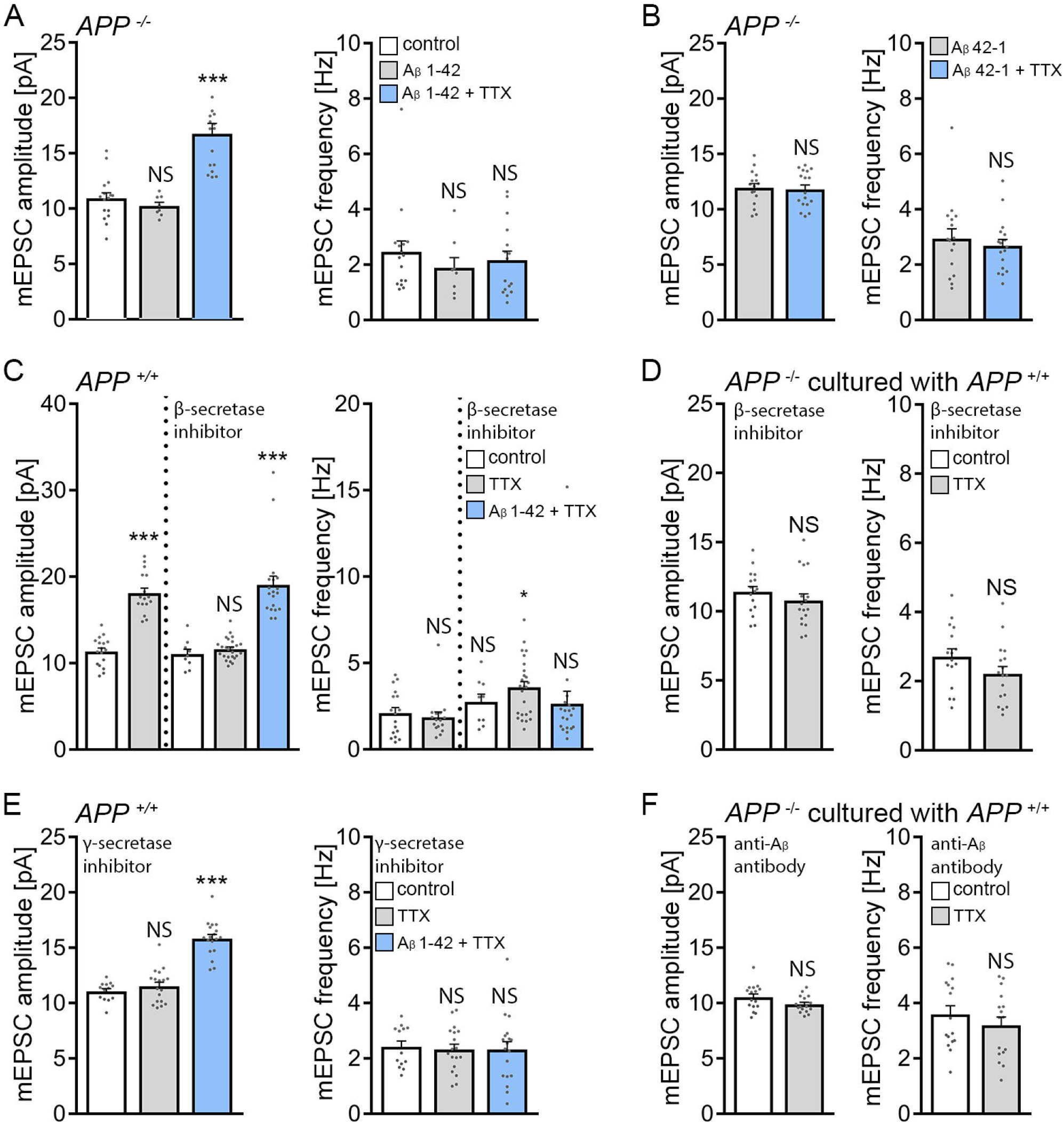
Amyloid-β_1 – 42_ rescues the ability of APP-deficient dentate granule cells to express homeostatic synaptic plasticity. **(A, B)** Group data of AMPA receptor mediated miniature excitatory postsynaptic currents (mEPSC) recorded from granule cells in vehicle-treated (control) and tetrodotoxin (TTX)- treated (2 μM, 2 d) *APP^−/−^* tissue cultures in the presence of amyloid-β1 – 42 (Aβ_1-42_; 1.5 μM) or Aβ_42-1_ (1.5 μM). (A: control, n = 16 cells from 4 cultures; Aβ_1-42_, n = 8 cells from 3 cultures; Aβ_1-42_ + TTX, n = 16 cells from 5 cultures; Kruskal-Wallis-test followed by Dunn’s post-hoc test; for mEPSC amplitudes one data point is outside the axis limits in the Aβ_1-42_ + TTX group; B: Aβ_42-1_, n = 16 cells from 4 cultures; Aβ_42-1_ + TTX, n = 17 cells from 5 cultures; Mann-Whitney test). **(C)** Group data of AMPA receptor mediated mEPSCs recorded from granule cells in vehicle-treated (control) and TTX-treated (2 μM, 2 d) wild-type tissue cultures. Pharmacological inhibition of the β-site APP cleaving enzyme (BACE inhibitor C3; 20 μM) prevents TTX-induced synaptic scaling; an effect that is reversed with Aβ_1-42_ (1.5. μM). Untreated: control, n = 17 cells from 5 cultures; TTX, n = 16 cells from 4 cultures. β-secretase inhibitor: control, n = 9 cells from 3 cultures; TTX, n = 25 cells from 7 cultures; Aβ_1-42_ + TTX, n = 19 cells from 5 cultures; Kruskal-Wallis-test followed by Dunn’s post-hoc test). **(D)** Group data of AMPA receptor mediated mEPSCs recorded from granule cells in vehicle-treated (control) and TTX-treated (2 μM, 2 d) *APP^−/−^* preparations co-cultured with *APP^+/+^* tissue (c.f., Figure 4). BACE inhibitor C3 (20 μM) prevents TTX-induced synaptic scaling (control, n = 16 cells from 5 cultures; TTX, n = 17 cells from 5 cultures; NS; Mann-Whitney test). **(E)** Pharmacological inhibition of γ – secretases with Begacestat (1 μM) prevents TTX-induced synaptic scaling of dentate granule cells in wild-type tissue cultures; an effect that is reversed with Aβ_1-42_ (1.5. μM). control, n = 13 cells from 4 cultures; TTX, n = 18 cells from 6 cultures; Aβ_1-42_ + TTX, n =18 cells from 6 cultures; Kruskal-Wallis-test followed by Dunn’s post-hoc test). (F) In the presence of a specific anti-Aβ antibody (1.3 μg/ml) TTX-induced synaptic scaling is not observed (results of the experiments in wild-type tissue reported in the main text; n = 16 cells from 6 cultures in each group; Mann-Whitney test). Values represent mean ± s.e.m. (* p < 0.05; *** p < 0.001; NS, no significant difference).

### Pharmacological inhibition of β-secretases in wild-type tissue cultures blocks homeostatic synaptic plasticity

Due to the fact that endogenous Aβ is produced by sequential cleavage of APP by β- and γ-secretases (Chow et al., 2010; Zheng and Koo, 2011), we next hypothesized that pharmacological inhibition of β-secretases (i.e., blocking the first step of the amyloidogenic processing pathway) should impede the ability of wild-type dentate granule cells to express TTX-induced synaptic scaling. Accordingly, wild-type cultures were treated with TTX (2 μM; 2 days) and with BACE inhibitor C3 [20 μM; 2 days;(Stachel et al., 2004)]. Pharmacological inhibition of β-secretases had no apparent effect on baseline mEPSC recordings, and a TTX-induced increase in mEPSC amplitudes was not observed. Exposure to Aβ_1-42_ (1.5 μM) together with BACE inhibitor C3 restored the ability of wild-type dentate granule cells to express homeostatic synaptic scaling (Figure 7*C*). Furthermore, in the presence of BACE inhibitor C3, TTX-induced synaptic scaling was not observed in *APP*^−^ preparations co-cultured with 5 *APP^+/+^* cultures on the same membrane insert (Figure 7*D*; c.f., Figure 4). These results suggest that APP/Aβ is part of an endogenous signaling pathway that mediates homeostatic synaptic plasticity.

### Pharmacological inhibition of γ-secretases in wild-type tissue cultures blocks homeostatic synaptic plasticity

BACE not only cleaves APP but also targets several other substrates in the nervous system (Barao et al., 2016). Although Aβ_1-42_ rescued homeostatic synaptic plasticity in BACE inhibitor C3-treated wild-type cultures, we decided to err on the side of caution. Accordingly, another series of experiments was carried out using the γ-secretase inhibitor Begacestat [GSI-953; 1 μM; (Martone et al., 2009)] to block the second enzymatic step of the amyloidogenic processing pathway. As shown in Figure 7*E*, a compensatory increase in mEPSC amplitudes was not observed in the presence of Begacestat after TTX treatment. Again, Aβ_1-42_ rescued the ability of wild-type dentate granule cells to express homeostatic synaptic plasticity in the presence of the γ-secretase inhibitor (Figure 7*E*). Consistent with these findings, Aβ sandwich-ELISA confirmed that endogenous Aβ_1-42_ is significantly reduced upon pharmacologic inhibition of β- or γ-secretases in our experimental setting [control, 115 ± 10 pg/ml; BACE inhibitor C3, 14 ± 0.9 pg/ml; Begacestat, 4 ± 0.1 pg/ml; medium harvested form n = 3 – 6 wells per group; p < 0.001; one-way ANOVA].

### Scavenging endogenous Aβ with a specific antibody blocks homeostatic synaptic plasticity in wild-type tissue cultures

Finally, we carried out experiments in wild-type cultures using an antibody that binds endogenous Aβ [anti-Aβ_11-28_; BNT77; 1.3 μg/ml; (Hashimoto et al., 2010)]. No significant differences in mEPSC properties between vehicle-only and TTX-treated wild-type dentate granule cells were observed in these experiments (control, 10.9 ± 0.4 pA; TTX 10.6 ± 0.3 pA n = 15 cells per group; p = 0.62; Mann-Whitney-test). Similarly, in the experimental setting in which *APP^−/−^* tissue was co-cultured with *APP^+/+^* cultures, again, no TTX-induced synaptic scaling was observed in the *APP^−/−^* cultures in the presence of the anti-Aβ antibody (Figure 7*F*; c.f., Figure 4).

Taken together, we conclude that APP is a key regulator of homeostatic synaptic plasticity and that Aβ-dependent signaling pathways account for the ability of cultured dentate granule cells to express TTX-induced homeostatic plasticity of excitatory synapses.

### Pharmacologic inhibition of NMDA-Rs rescues homeostatic synaptic plasticity in APP-deficient preparations

What are the downstream signaling pathways of APP/Aβ mediated homeostatic synaptic plasticity? Previous work revealed that Aβ affects Hebbian plasticity via modulation of *N*-methyl-D-aspartate receptors [NMDA-R; (Chen et al., 2002; Snyder et al., 2005; Zhang et al., 2009)]. Considering the role of NMDA-R in homeostatic synaptic plasticity [i.e., NMDA-R inhibition triggers synaptic up-scaling; (Sutton et al., 2006)], we tested whether pharmacologic inhibition of NMDA-R rescues homeostatic synaptic plasticity in the absence of Aβ (i.e., in *APP^−/−^* tissue cultures). In the presence of the NMDA-R antagonist D-APV (50 μM), a significant increase in mEPSC amplitudes was noted in TTX-treated APP^−/−^ granule cells (Figure 8*A, B*). Interestingly, recordings of NMDA-R-mediated mEPSCs showed no significant differences between APP^−/−^ and APP^+/+^ granule cells (APP^+/+^, 17.6 ± 0.7 pA; APP^−/−^ 17.0 ± 0.6 pA; n = 9 and 10 cells, respectively; p = 0.66; Mann-Whitney test), confirming once more that basic functional differences between the two genotypes do not trivially explain our major findings (c.f., Figures 2 and 3). These results indicate that Aβ acts on intracellular Ca^2+^ sensors or effectors, which respond to the TTX-induced reduction of intracellular Ca^2+^ in our experiments.

**Figure 8:**
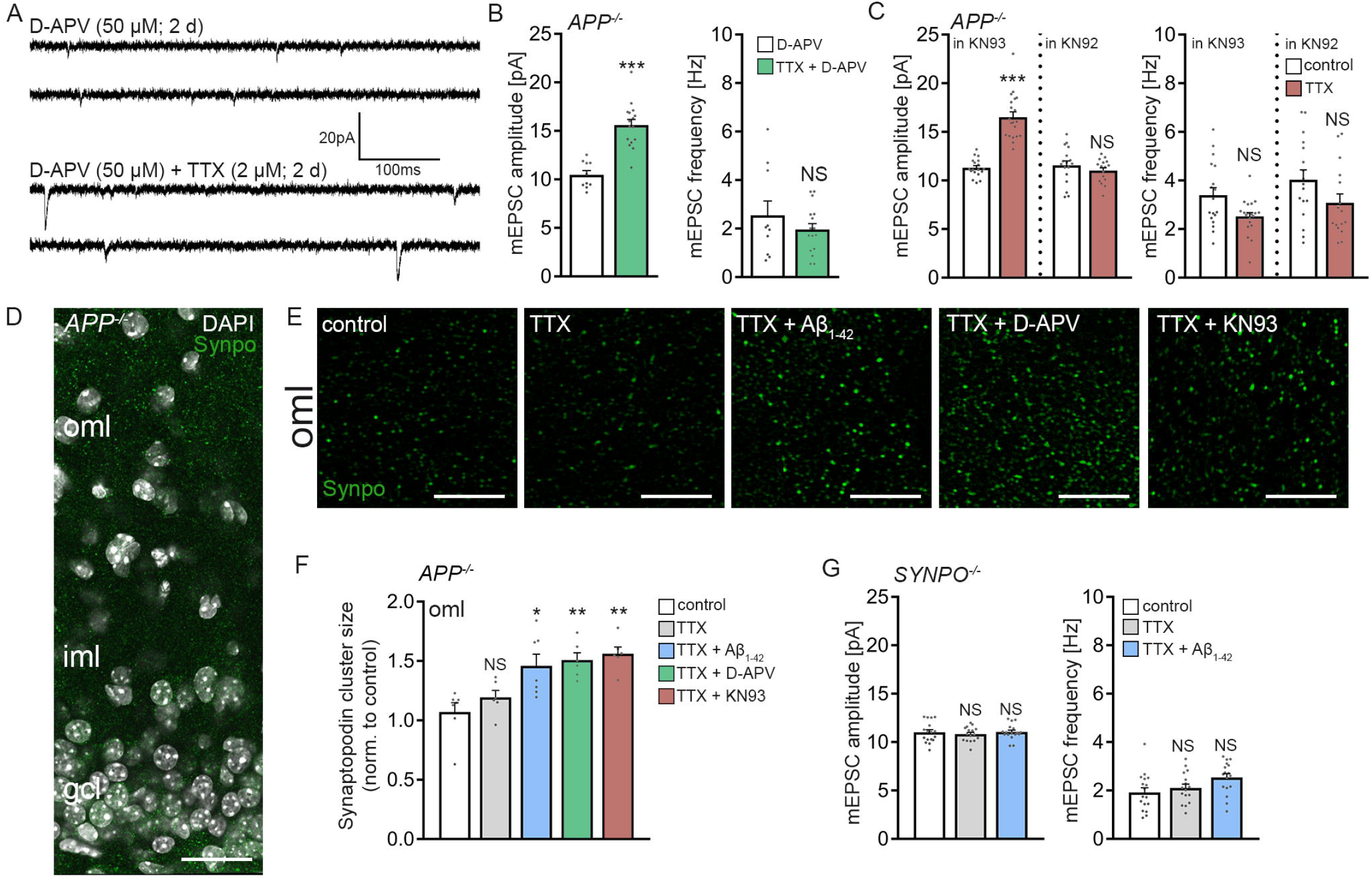
Role of Ca^2+^ signaling and synaptopodin in APP/amyloid-β mediated homeostatic synaptic scaling. **(A, B)** Sample traces and group data of AMPA receptor mediated miniature excitatory postsynaptic current (mEPSCs) recorded from granule cells in D-APV (50 μM; 2 d) treated and D-APV (50 μM) + TTX-treated (2 μM, 2 d) *APP^−/−^* tissue cultures (D-APV, n = 10 cells from 3 cultures; TTX + D-APV, n = 16 cells from 4 cultures; Mann-Whitney test). **(C)** Group data of AMPA receptor mediated mEPSCs from granule cells in *APP^−/−^* cultures recorded in the presence of the calcium/calmodulin-dependent protein kinase II (CamKII) inhibitor KN93 or the inactive analogue KN92 (KN93: control, n = 18 cells from 5 cultures; TTX = 20 cells from 5 cultures; KN92: control, n = 17 cells from 5 cultures; TTX = 16 cells from 5 cultures; one data point in mEPSC frequency outside the axis limits; NS; Mann-Whitney test). **(D)** Example of a *APP^−/−^* tissue cultures immunostained for synaptopodin (Synpo, green). Cluster sizes were assessed in the outer molecular layer (oml) of the dentate gyrus. (iml, inner molecular layer; gcl, granule cell layer; DAPI, nuclear stain). Scale bar = 50 μm. **(E)** Examples of analysed visual fields (six fields per culture and condition) at higher magnification. All treatments 2 days. TTX, 2μM; Aβ_1-42_, 1.5 μM; D-APV, 50 μM, KN93, 5 μM. Scale bar 5 μm **(F)** Group data of synaptopodin cluster sizes in the respective groups. Values normalized to controls (control, n = 7 cultures; TTX, n = 6 cultures; TTX + Aβ_1-42_, n = 7 cultures; TTX + D-APV, n = 6 cultures; TTX + KN93, n = 6 cultures; Kruskal-Wallis-test followed by Dunn’s post-hoc test). **(G)** Group data of AMPA receptor mediated mEPSCs recorded from granule cells in synaptopodin-deficient (*SYNPO*^−/−^) tissue cultures. Aβ_1-42_ (1.5 μM, 2 d) does not rescue TTX-induced homeostatic synaptic plasticity (untreated, n = 17 cells from 6 culltures; TTX, n =16 cells from 5 cultures; TTX + Aβ_1-42_, n =17 cells from 5 cultures; Kruskal-Wallis-test followed by Dunn’s post-hoc test). Values represent mean ± s.e.m. (* p < 0.05; ** p < 0.01; *** p < 0.001; NS, no significant difference).

### Pharmacologic inhibition of CamKII rescues homeostatic synaptic plasticity in APP-deficient preparations

Aβ has been linked to inactivation of calcium/calmodulin-dependent protein kinase II [CamKII; (Gu et al., 2009; Townsend et al., 2007; Zhao et al., 2004); but see (Opazo et al., 2018)]. Hence, we reasoned that in the absence of APP/Aβ, pharmacological inhibition of CamKII might also rescue TTX-induced synaptic scaling. Indeed, increased mEPSC amplitudes were observed in *APP^−/−^* cultures upon TTX treatment in the presence of KN-93 (5 μM), but not with the inactive analogue KN-92 (5 μM; Figure 8*C*). Hence, NMDA-R-mediated CaMKII-dependent downstream signaling pathways seem to be involved in APP/Aβ-mediated homeostatic synaptic plasticity.

### Role of synaptopodin in APP/Aβ mediated homeostatic synaptic plasticity

In previous work, we demonstrated that the actin-binding molecule synaptopodin, which is a marker and essential component of the Ca^2+^-storing spine apparatus organelle (Deller et al., 2003), is required for the expression of homeostatic synaptic plasticity (Vlachos et al., 2013). In this context, a Ca^2+^-dependent compensatory increase in synaptopodin clusters was observed, consistent with a negative feedback mechanism that mediates homeostatic synaptic plasticity (Vlachos et al., 2013). Notably, an association between synaptopodin and peptides corresponding to CamKIIα and CamKIIβ has been recently reported (Konietzny et al., 2019).

To test whether activity-dependent changes of synaptopodin are part of a negative feedback mechanism that requires APP/Aβ-dependent signalling, *APP^−/−^* cultures were once again treated with TTX (2 μM; 2 d). Immunostained synaptopodin clusters were assessed in the outer molecular layer of the dentate gyrus, and the previously reported compensatory increase in synaptopodin cluster sizes was not observed in TTX-treated *APP^−/−^* cultures [c.f., (Vlachos et al., 2013)]. We then tested whether in the presence of Aβ_1-42_, TTX-induced changes in synaptopodin clusters are triggered. Indeed, the same protocol that rescues homeostatic synaptic plasticity in *APP^−/−^* preparations also induces synaptopodin changes similar to that observed in wild-type cultures (Figure 8*F*).

Does pharmacological inhibition of NMDA-Rs and CamKII rescue homeostatic synaptic scaling-associated changes in synaptopodin cluster properties of *APP^−/−^* cultures? As shown in Figure 8F, in the presence of D-APV (50 μM) or KN93 (5 μM), a significant increase in synaptopodin clusters was observed in response to TTX (2 μM, 2 days). These results indicate that APP/Aβ is part of a Ca^2+^-dependent negative feedback mechanism that regulates synaptopodin cluster properties in an NMDA-R- and CamKII-dependent manner.

Finally, based on the observation that Aβ_1-42_ rescues TTX-induced homeostatic synaptic plasticity, we tested for the effects of Aβ_1-42_ in synaptopodin-deficient dentate granule cells, which do not form spine apparatus organelles and show defects in homeostatic synaptic plasticity (Vlachos et al., 2013). Whereas the previously reported deficit in TTX-induced homeostatic synaptic plasticity was reproduced in these experiments, Aβ_1-42_ did not rescue homeostatic synaptic plasticity in *SYNPO*^−/−^ dentate granule cells (Figure 8*G*). Taken together, we propose that synaptopodin is one of the downstream molecular targets required for APP/Aβ-mediated homeostatic synaptic plasticity.

## DISCUSSION

We regard the significant finding of this study to be the discovery of a previously unknown APP/Aβ phenotype that becomes salient after network activity is altered (i.e., blocked by TTX for a prolonged period of time). Under these conditions, *APP^−/−^* dentate granule cells do not scale their excitatory synapses in a homeostatic manner. Surprisingly, this plasticity phenotype does not depend on APPsα, which is generated from APP via non-amyloidogenic processing and which has been linked to Hebbian plasticity, but on Aβ, which is generated from APP via processing along the amyloidogenic pathway. Consistent with this observation, pharmacologic inhibition of β- and γ-secretases as well as scavenging endogenous Aβ with antibodies blocked TTX-induced synaptic scaling in wild-type tissue cultures. Aβ-dependent synaptic scaling required modulation of downstream Ca^2+^-dependent signaling pathways, including NMDA-Rs and CamKII, as well as synaptopodin—a molecule essential for the formation of the Ca^2+^-storing spine apparatus. Together, these results reveal a direct involvement of amyloidogenic APP processing and Aβ in homeostatic synaptic plasticity. This involvement raises the intriguing possibility that changes in the balance of APP processing along the two major APP-processing pathways could also lead to changes in the balance of Hebbian and homeostatic plasticity in the brain (Figure 9).

**Figure 9:**
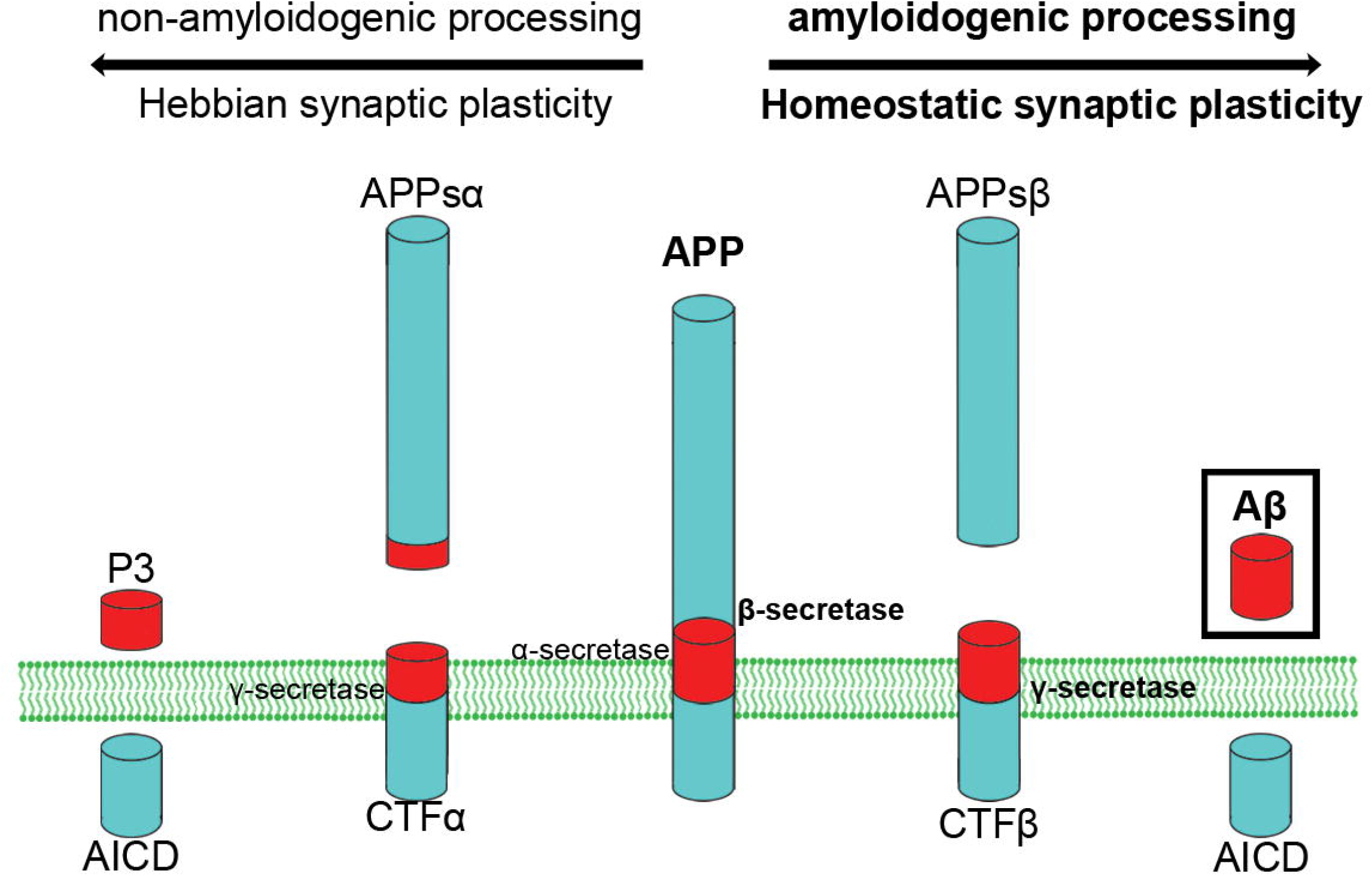
Possible implications for APP processing in synaptic plasticity. A firm link between the non-amyloidogenic processing pathway (i.e., APP secreted ectodomain APPsα and the ability of neurons to express long-term potentiation of excitatory neurotransmission has been established. The results of the present study show that the amyloidogenic processing pathway, which produces amyloid-β (Aβ) may serve homeostatic synaptic plasticity. It is thus interesting to speculate, that signaling pathways that influence APP processing via β-secretase or γ-secretases may set a balance between Hebbian and homeostatic synaptic plasticity in neural networks

Studies employing mouse mutants lacking APP or APP-gene family members have established a firm link between APP and the ability of neurons to express synaptic plasticity (Müller et al., 2017). Specifically, the ability of neurons to express LTP of excitatory neurotransmission is impaired in aged *APP^−/−^* mice [e.g., (Dawson et al., 1999; Ring et al., 2007; Seabrook et al., 1999; Tyan et al., 2012)]. APPsα could rescue this deficit [e.g., (Fol et al., 2016; Hick et al., 2015; Richter et al., 2018; Ring et al., 2007) using recombinant, viral, and genetic strategies, suggesting a “plasticity-promoting” role of APP via the non-amyloidogenic processing pathway (Mockett et al., 2017). Based on these observations, we assumed that another major form of synaptic plasticity [i.e., homeostatic synaptic plasticity (Turrigiano et al., 1998)], could also depend on APP or one of its diverse processing products. Indeed, *APP^−/−^* neurons failed to scale their synapses after TTX treatment, implicating APP in homeostatic plasticity, but APPsα could not rescue this APP-dependent plasticity phenotype. Similarly, scavenging endogenous APPsα or treatment with recombinant APPsα did not affect TTX-induced homeostatic synaptic plasticity in wild-type dentate granule cells. This phenomenon cannot be trivially explained by an inability of dentate granule cells to respond to APPsα because the exogenous application of APPsα robustly enhances LTP in these neurons (Taylor et al., 2008). Together, these observations suggest that the plasticity-promoting effect of APPsα (Mockett et al., 2017) should be narrowed down and specified: APP processing via the non-amyloidogenic pathway promotes the ability of neurons to express LTP, but not necessarily all forms of synaptic plasticity because our data demonstrates that APPsα is not a key regulator of homeostatic synaptic plasticity.

The inability to rescue homeostatic plasticity via APPsα raised the intriguing possibility that APP processing via the “amyloidogenic pathway” could serve homeostatic synaptic plasticity. In the presence of Aβ, the homeostatic synaptic response to TTX was fully restored in APP^−/−^ preparations. Moreover, we report that pharmacologic inhibition of β-secretases blocks TTX-induced homeostatic synaptic plasticity in wild-type cultures. Because pharmacologic inhibition of γ-secretase had a similar effect, and Aβ rescued homeostatic synaptic plasticity in the presence of β- or γ-secretase inhibitors, other APP cleavage products [e.g., APPsβ (or additional factors arising from β- or γ-secretase processing)] are unlikely to account for our significant findings. In a previous study, however, pharmacologic inhibition of γ-secretases did not block TTX-induced homeostatic synaptic plasticity (Pratt et al., 2011). These earlier experiments were carried out in primary hippocampal neurons rather than in organotypic tissue cultures, and L685,458 (5 μM) was used instead to Begacestat (1 μM) in our study. These differences may explain the inconsistent results. Notably, we confirmed that pharmacologic inhibition of γ-secretase with Begacestat reduced endogenous Aβ levels in our experimental setting, and exogenous Aβ_1-42_ rescued homeostatic synaptic plasticity in Begacestat-treated tissue cultures.

To provide further evidence for the role of Aβ in neuronal physiology, we employed antibodies against Aβ and scavenged endogenously produced Aβ from wild-type tissue cultures. This approach abolished homeostatic synaptic scaling in the wild-type cultures and prevented synaptic scaling in *APP^−/−^* preparations cultured together with *APP^+/+^* tissue; whereas a specific antibody against APPsα did not affect TTX-induced synaptic scaling. Together, we propose a model in which APP processing via the non-amyloidogenic pathway and the amyloidogenic pathway influences the ability of a neuron to express distinct forms of synaptic plasticity: processing along the non-amyloidogenic pathway may promote Hebbian plasticity (e.g., LTP), whereas processing along the amyloidogenic pathway—or increased availability of Aβ (Gilbert et al., 2016)—promotes homeostatic synaptic plasticity (Figure 9).

What are the downstream pathways activated by Aβ in the context of homeostatic plasticity? Consistent with previous work on Hebbian plasticity (Gu et al., 2009; Sinnen et al., 2016; Snyder et al., 2005; Zhang et al., 2009; Zhao et al., 2004), we could link the homeostatic plasticity effects of Aβ to NMDA-Rs and CamKII. Considering the opposing roles of NMDA-R and CamKII in Hebbian and homeostatic synaptic plasticity, we speculated that Aβ-signaling could be part of a Ca^2+^-dependent negative feedback mechanism that mediates homeostasis. Indeed, pharmacologic inhibition of NMDA-Rs and CamKII rescued homeostatic synaptic plasticity in *APP^−/−^* preparations, suggesting that Aβ acts on intracellular Ca^2+^ sensors or effectors, which responded to the TTX-induced reduction in intracellular Ca^2+^ in our experiments. Consistent with this suggestion, the previously reported Ca^2+^-dependent compensatory adjustment of synaptopodin was not observed in *APP^−/−^* preparations [c.f., (Vlachos et al., 2013)], and interventions that rescued TTX-induced synaptic scaling in *APP^−/−^* dentate granule cells also triggered compensatory changes in synaptopodin. Moreover, Aβ was not able to rescue homeostatic synaptic plasticity in *SYNPO^−/−^* preparations, demonstrating that synaptopodin is an essential downstream target through which APP/Aβ asserts its effects on homeostatic synaptic plasticity.

In previous work, we have shown that the plasticity-related protein synaptopodin controls the ability of neurons to express both Hebbian and homeostatic synaptic plasticity (Deller et al., 2003; Jedlicka et al., 2009; Vlachos et al., 2013; Vlachos et al., 2009). Animals lacking this protein do not form spine apparatus organelles and exhibit deficits in Hebbian and homeostatic synaptic plasticity (Deller et al., 2003; Jedlicka et al., 2009; Vlachos et al., 2013) because the Ca^2+^-dependent accumulation of AMPA-Rs at excitatory postsynapses is impaired [e.g., (Maggio and Vlachos, 2018; Vlachos et al., 2013; Vlachos et al., 2009)]. The results of the present study call for a systematic assessment of the activity-dependent molecular pathways through which APP processing via the “amyloidogenic/homeostatic”‘ and “non-amyloidogenic/Hebbian plasticity promoting” pathways affect synaptopodin-mediated synaptic plasticity. It may be important to mention in this context that changes in synaptopodin expression have been reported in brain tissue from AD subjects (Reddy et al., 2005) and synaptopodin has been recently linked to autophagy of phospho-MAPT/Tau (Ji et al., 2019). Moreover, a reduction in synaptopodin expression seems to ameliorate symptoms in a transgenic mouse model of AD (Aloni et al., 2019).

A role of homeostatic synaptic plasticity in AD has been recently discussed (Jang and Chung, 2016; Styr and Slutsky, 2018). In the context of our data, which suggest a physiological role of Aβ, it can be speculated that the previously reported “synaptotoxic” effects of Aβ [e.g., (Mucke and Selkoe, 2012; Westmark, 2013; Zott et al., 2019)] are the result of a pathological over-activation of molecular machinery promoting synaptic homeostasis under physiological conditions. Consistent with this pathophysiological concept, Aβ mediated “alterations” in LTP in AD models might reflect –at least in part– enhanced homeostatic synaptic plasticity, which rapidly returns potentiated synapses to baseline upon LTP induction. Clearly, the biological consequences of the “amyloidogenic/homeostatic” and the ‘”non-amyloidogenic/Hebbian plasticity promoting” pathways warrant further investigation in the AD context, particularly because the amyloidogenic pathway is an important target for therapeutic intervention in AD (Coimbra et al., 2018; Egan et al., 2019; Yan and Vassar, 2014). Hence, some of the side-effects observed in patients treated with β-secretase inhibitors (Coimbra et al., 2018; Egan et al., 2019) may have been caused by interference with the ability of healthy neurons to express homeostatic synaptic plasticity.

## MATERIALS AND METHODS

### Ethics statement

Mice were maintained in a 12 h light/dark cycle with food and water available ad libitum. Every effort was made to minimize distress and pain of animals. All experimental procedures were performed according to German animal welfare legislation and approved by the appropriate animal welfare committee and the animal welfare officer of Freiburg University.

### Animals

Wild-type *C57BL/6J, SYNPO^−/−^* (Deller et al., 2003), *APP^−/−^* (Li et al., 1996), *APPsα-KI* mice (Ring et al., 2007) and their wild-type littermates were used in this study.

### Preparation of tissue cultures

Enthorhino-hippocampal tissue cultures were prepared at postnatal day 4-5 from mice of either sex as previously described (Vlachos et al. 2012). The incubation medium consisted of 50 % (v/v) minimum essential medium (MEM), 25 % (v/v) basal medium eagle (BME), 25 % (v/v) heat inactivated normal horse serum (NHS), 25 mM HEPES, 0.15 % (w/v) NaHCO3, 0.65 % (w/v) Glucose, 0.1 mg/ml Streptomycin, 100U/ ml Penicillin and 2 mM Glutamax. The pH was adjusted to 7.3 and the medium was changed 3 times per week. Tissue cultures were allowed to mature *in vitro* for at least 18 days before any experimental procedure.

### Whole cell patch-clamp

Whole cell patch-clamp recordings of dentate gyrus granule cells were carried out at 35°C. The bath solution contained 126 mM NaCl, 2.5 mM KCl, 26 mM NaHCO_3_, 1.25 mM NaH_2_PO_4_, 2 mM CaCl_2_, 2 mM MgCl_2_ and 10 mM glucose and was saturated with 95% O2 / 5% CO2. For miniature and spontaneous AMPA receptor mediated post synaptic current (m/sEPSC) recordings as well as current clamp (input-output) recordings patch pipettes contained 126 mM K-gluconate, 4 mM KCl, 4 mM ATP-Mg, 0.3 mM GTP-Na2, 10 mM PO-Creatine, 10 mM HEPES and 0.1% Biocytin (pH = 7.25 with KOH, 290 mOsm with sucrose). For NMDA receptor mediated mEPSC recordings patch pipettes contained 120 mM CsCH3SO3, 8 mM CsCl, 1 mM MgCl2, 0.4 mM EGTA, 2 mM ATP-Mg, 0.3 mM GTP-Na2, 10 mM PO-Creatine, 10mM HEPES and 5 mM QX-314 (pH = 7.25 with CsOH, 295 mOsm with sucrose). For miniature and spontaneous inhibitory post synaptic current (m/sIPSC) recordings patch pipettes contained 40 mM CsCl, 90 mM K-gluconate, 1.8 mM NaCl, 1.7 mM MgCl2, 3.5 mM KCl, 0.05 mM EGTA, 2 mM ATP-Mg, 0.4 mM GTP-Na2, 10 mM PO-Creatine, 10 mM HEPES (pH = 7.25 with CsOH, 270 mOsm with sucrose). AMPA receptor – mediated mEPSCs were recorded in the presence of 10 μM D-APV and 0.5 μM TTX, NMDA-R-mediated mEPSCs in the presence of 10 μM CNQX and 0.5 μM TTX and mIPSCs in the presence of 0.5 μM TTX, 10 μM D-APV and 10 μM CNQX. sEPSCs were recorded without the addition of any drugs in the bath solution and sIPSCs in the presence of 10 μM D-APV and 10 μM CNQX. Current clamp recordings were performed in the presence of 10 μM D-APV, 10 μM CNQX and 10 μM Bicuculline methiodide. Neurons were recorded at a holding potential of −70 mV for m/sIPSCs and AMPA receptor mediated m/sEPSCs. NMDA receptor mediated mEPSCs were acquired at +40 mV. For current clamp recordings neurons were hyperpolarized at −100 pA and then depolarized up to +200 pA with one-second-long 10 pA current injection steps. Series resistance was monitored in 2 min intervals, and recordings were discarded if the series resistance reached ≥30 MΩ and the leak current changed significantly.

### Reconstruction of dendritic trees and spine density analysis

Dentate granule cells were patched with Alexa 568 (10 μM) added to the internal solution and filled for 10 minutes in whole-cell configuration for visualization of identified cells. Confocal image stacks of granule cells (512×512 pixel, voxel size 0.49 x 0.49 x 2μm) were acquired directly at the electrophysiology setup using a Zeiss LSM Exciter confocal microscope with a 40x water immersion objective lens (0.8NA; Zeiss). Granule cells were manually reconstructed in 3D and analyzed using Neurolucida/NeuroExplorer software (MBF Bioscience). Total dendritic length (TDL) was calculated as the sum of length of all reconstructed dendritic segments of a given cell. For Sholl analysis, concentric spheres with diameters increasing in 20 μm increments were drawn around the cell soma, and the number of dendrites intersecting each sphere was calculated. Outer molecular layer (OML) segments were imaged with higher scan zoom and spine densities were determined as described previously (Hick et al., 2015).

### Immunostaining and imaging

Tissue cultures were fixed in a solution of 4 % (w/v) paraformaldehyde (PFA) and 4 % (w/v) sucrose in 0.01 M phosphate-buffered saline (PBS) for 1 h, followed by 2 % (w/v) PFA and 30 % (w/v) sucrose in 0.01 M PBS overnight. For synaptopodin staining. 30 μm cryo-sections were prepared, and stained with antibodies against synaptopodin (1:1000; SE-19 Sigma-Aldrich, RRID: AB_261570). Sections were incubated for 1 h with 10 % (v/v) normal goat serum (NGS) in 0.5 % (v/v) Triton X-100-containing PBS to reduce unspecific antibody binding and incubated for 24 h at 4 °C with the primary antibody in PBS with 10% NGS and 0.1 % Triton X-100. Sections were washed and incubated for 4 h with appropriate Alexa-labelled secondary antibodies (Invitrogen; 1:1000, in PBS with 10 % NGS, 0.1 % Triton X-100). DAPI nuclear stain was used to visualize cytoarchitecture (1:5000; in 0.01 M PBS for 15 min). Sections were washed with 0.1 M PBS, transferred onto glass slides and mounted for visualization with anti-fading mounting medium. Confocal images were acquired using a Leica TCS SP8 laser scanning microscope with 20x (NA 0.75; Leica), 40x (NA 1.30; Leica) and 63x (NA 1.40; Leica) oil-submersion objectives.

### *Post-hoc* identification of recorded neurons

Tissue cultures were fixed in a solution of 4 % (w/v) and 4 % (w/v) sucrose in 0.01 M PBS for 1 h. The fixed tissue was incubated for 1 h with 10 % (v/v) NGS and 0.5 % (v/v) Triton X-100 in 0.01 M PBS. Biocytin (Sigma-Aldrich Cat# B4261) filled cells were stained with Alexa-488, −568 or −633 conjugated Streptavidin (Thermo Fisher Scientific; 1:1000; in 0.01 M PBS with 10 % NGS and 0.1 % Triton X100) for 4 h and DAPI (Thermo Fisher Scientific) staining was used to visualize cytoarchitecture (1:5000; in 0.01 M PBS for 15 min). Slices were washed, transferred and mounted onto glass slides for visualization. Virus expressing and streptavidin stained granule cells were visualized with a Leica TCS SP8 laser scanning microscope with 20x (NA 0.75; Leica), 40x (NA 1.30; Leica) and 63x (NA 1.40; Leica) oil-submersion objectives.

### Electron microscopy

Tissue cultures were fixed and prepared as described previously (Vlachos et al., 2013). Briefly, cultures were fixed in 15% picric acid, 4% glutaraldehyd, 4% PFA in PB-buffer. After washing in PB, cultures were osmicated (0.5% O_s_O_4_ in PB, 30 minutes), dehydrated (70% ethanol containing 1% uranyl acetate), and embedded between liquid release-coated slides and coverslips. Cultures were re-embedded in blocks and serial ultrathin sections were collected on single-slot Formvar-coated copper grids. Ultrathin sections were examined in a Zeiss electron microscope (EM 109).

### Pharmacology

Tissue cultures were treated with tetrodotoxin (TTX; 2 μM; BioTrend, Cat# BN0518), D-APV (50 μM; Tocris, Cat# 0106) synthetic Aβ_1-42_ (1.5 μM; Bachem, Cat# 4014447) or Aβ_42-1_ (1.5 μM; Bachem, Cat# H-3976) (Novotny et al., 2016), recombinant APPsα (10 nM) (Hick et al., 2015), BACE inhibitor C3 (20 μM; Calbiochem, Sigma-Aldrich Cat# 565788), Begacestat (1 μM; Tocris, Cat# 4283), KN-92 (5 μM; Tocris, Cat# 4130), KN-93 (5 μM; Tocris, Cat# 1278), BNT77 (1.3 μg/ml; Wako, Cat# 014-26881, RRID: AB_2827702), JRD32 [1.3 μg/ml; (Hick et al., 2015), RRID: AB_2827703] for 2 days. For the Aβ treatment, PBS was added to the lyophilized Aβ peptides, followed by 3 x 1 min vortexing on wet ice and stocks were kept at −80°C. Before treating the tissue cultures, Aβ_1-42_ and Aβ_42-1_ were slowly thawed in ice and vortexed again before adding in the medium.

### Viral transduction

Tissue cultures were transfected between 3–4 div by adding 1 μl of AAV-Syn/Venus_T2A_APPsα (AAV-APPsα) or 1 μl of AAV-Syn/APP695_T2A_Venus (AAV-flAPP-Venus; kindly provided by Drs. Christian Buchholz and Tobias Abel) directly on top of each culture. Tissue cultures were then left to mature at least until 18 div before experimental assessment.

### ELISA assay and Aβ peptide quantification

Incubation medium from vehicle-only- (control) and pharmacologically-treated 3-week-old tissue cultures was collected, frozen immediately with dry ice, and stored at −80°C (vehicle, n = 6 wells with 36 tissue cultures; BACE inhibitor C3, n = 3 wells with 18 tissue cultures; Begacestat, n = 3 wells with 18 tissue cultures). For the detection of Aβ_1-42_ a V-PLEX™ Aβ Peptide Panel 1 Kit (MesoScale Discovery, Rockville, USA, Cat# K15199E, RRID: AB_2827747) was used with a ruthenylated anti-Aβ antibody (4G8 clone). The Aβ peptide concentration was determined using the MSD Discovery Workbench software (MesoScale Discovery, Rockville, USA).

### Quantification and statistics

Analyses were performed with the person analyzing the data blind to experimental condition. One to two tissue cultures were used from each animal. Electrophysiological data were analyzed using pClamp 10.7 software suite (Molecular Devices), MiniAnalysis (Synaptosoft) and Igor Pro 7 (Wavemetrics). 150-300 events were analyzed per recorded neuron. Sizes of immunolabelled synaptopodin clusters were assessed using the FIJI – ImageJ software package (available from https://imagej.net/ImageJ) as described previously (Vlachos et al., 2013). Statistical comparisons were made using Mann-Whitney test (to compare two groups), Kruskal-Wallis-test followed by Dunn’s *post-hoc* test for multiple group testing or one or two-way ANOVA as indicated in the figure captions and text (GraphPad Prism 7, GraphPad software, USA). P-values of less than 0.05 were considered a significant difference. All values represent mean ± standard error of the mean. In the figures * denotes p < 0.05, ** p < 0.01 and *** p < 0.001; no significant differences are indicated with ‘NS’.

### Digital illustrations

Confocal image stacks were exported as 2D-projections and stored as TIF files. Figures were prepared using Photoshop graphics software (Adobe, San Jose, CA, USA). Image brightness and contrast were adjusted.

## ACKNOWLEDGEMENTS

We thank Anke Biczysko, Sussana Glaser, Katrin Moschke, and Barbara Joch for excellent technical assistance. We also thank Drs. Christian Buchholz and Tobias Abel for providing AAV vectors. The work was supported by the Federal Ministry of Education and Research, Germany (BMBF ‘OGEAM’ to T.D and U.M.), under Germany’s Excellence Strategy within the framework of the Munich Cluster for Systems Neurology (EXC 2145 SyNergy– ID 390857198, to SFL) and by Deutsche Forschungsgemeinschaft (MU1457/14-1 to U.M., FOR 1332 to U.M., T.D., A.V., CRC1080 T.D. and A.V., and CRC 974. to A.V.).

## AUTHOR CONTRIBUTIONS

The study was conceived and supervised by AV. Experiments were designed by CG, UM, TD and AV. CG, MF and DB performed experiments and analyzed the data. Sandwich-ELISA experiments were supervised by SFL. CB and UM provided recombinant APPsα, anti-APPsα antibodies, APP^−/−^ and APPsα-KI mice. The manuscript was written by AV with the help of CG, SFL, UM and TD. All authors were involved in data interpretation and critically revising the manuscript.

