## Supplementary material for "Amyloid-beta mediates homeostatic synaptic plasticity": Figure S1

### SUPPLEMENTAL INFORMATION

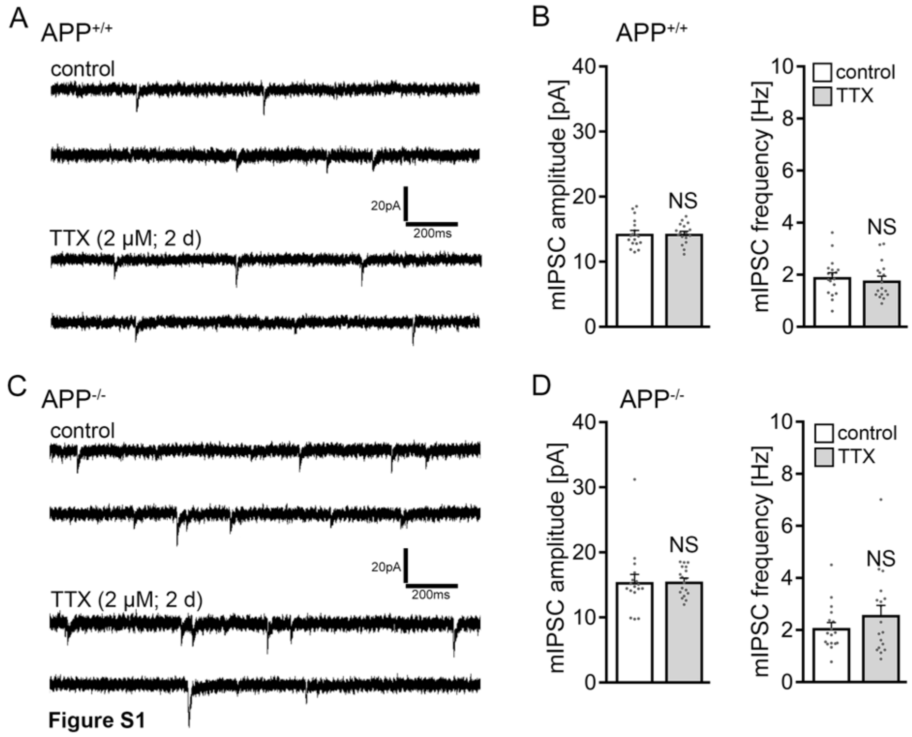


**Figure S1: Dentate granule cells do not adjust inhibitory synapses following TTX treatment**
**(A, B)** Sample traces and group data of miniature inhibitory postsynaptic currents (mIPSCs) recorded from granule cells in vehicle-only-treated (control) and tetrodotoxin (TTX)-treated $APP^{+/+}$ cultures (control, n = 18 cells from 6 cultures; TTX, n = 17 cells from 6 cultures; Mann-Whitney test). **(C, D)** Sample traces and group data of APP-deficient ($APP^{-/-}$) dentate granule cells (control, n = 17 cells from 6 cultures; TTX, n = 18 cells from 6 cultures; Mann-Whitney test). Values represent mean ± s.e.m. (NS, no significant difference).
